# Antibiotic-induced microbiota depletion impairs the pro-regenerative response to a biological scaffold in mice

**DOI:** 10.1101/2025.06.23.661121

**Authors:** Natalie Rutkowski, Brenda Yang, Elise Gray-Gaillard, Anna Ruta, Joscelyn C. Mejías, Michael Patatanian, Christopher Cherry, Katlin B. Stivers, Shri Ramanujam, Nathan L. Price, Franck Housseau, Drew M. Pardoll, Cynthia L. Sears, Jennifer H. Elisseeff

**Affiliations:** Translational Tissue Engineering Center, Johns Hopkins University, Baltimore, MD 21231; Translational Gerontology Branch, National Institute on Aging, Baltimore, MD 21224; Department of Medicine, Johns Hopkins University School of Medicine, Baltimore, MD 21287; Department of Oncology and Sidney Kimmel Comprehensive Cancer Center, Johns Hopkins University School of Medicine, Baltimore, MD 21287; Bloomberg∼Kimmel Institute for Cancer Immunotherapy, Johns Hopkins University School of Medicine, Baltimore, MD 21287; Department of Molecular Microbiology and Immunology, Johns Hopkins University Bloomberg School of Public Health, Baltimore, MD 21287

**Author notes:** **Author Contributions:** N.R., B.Y., E.G.G., A.R., F.H., D.M.P., C.L.S., and J.H.E designed research, N.R., B.Y., E.G.G., A.R., J.C.M., K.B.S., and S.R. performed research, N.R., B.Y., M.P., C.C., and J.H.E analyzed data, N.L.P. contributed new reagents/analytic tools, and N.R., B.Y., and J.H.E wrote the paper. **Competing Interest Statement:** J.H.E. holds equity in Unity Biotechnology and Aegeria Soft Tissue and is a consultant for Tessara. D.M.P. is consultant at Aduro Biotech, Amgen, Astra Zeneca, Bayer, Compugen, DNAtrix, Dynavax Technologies Corporation, Ervaxx, FLX Bio, Immunomic, Janssen, Merck, and Rock Springs Capital. D.M.P. holds equity in Aduro Biotech, DNAtrix, Ervaxx, Five Prime therapeutics, Immunomic, Potenza, Trieza Therapeutics. D.M.P. is a member of the scientific advisory board for Bristol Myers Squibb, Camden Nexus II, Five Prime Therapeutics, and WindMil. D.M.P. is a member of board of directors in Dracen Pharmaceuticals.

## Abstract

Therapeutic biological scaffolds promote tissue repair primarily through the induction of type 2 immunity. However, systemic immunological factors–including aging, sex, and previous infections–can modulate this response. The gut microbiota is a well-established modulator of immune function across organ systems, yet its influence on type 2-mediated repair remains underexplored. Here, we establish a bidirectional relationship between the gut microbiota and biological scaffold-mediated tissue repair. Utilizing a conventionalized germ-free mouse, we demonstrate that scaffold implantation induces compositional and functional changes in the gut microbiome, particularly affecting amino acid biosynthesis. Additionally, in a model of antibiotic-induced microbiota depletion (AIMD), we show that dysbiosis disrupts key immune regulators of type 2 immunity, including reductions in eosinophils, pro-regenerative macrophages, and IL-4-producing CD4^+^ T cells. At 6 weeks post-scaffold implantation, we observed a significant decrease in myocytes with centrally located nuclei alongside an upregulation in pro-fibrotic gene expression with antibiotic treatment. These findings provide insights into the influence of the gut microbiota on type 2-mediated tissue repair.

**Significance Statement:** Antibiotics are routinely administered perioperatively to prevent infection during surgeries and biomaterial implantation. Here, we demonstrate that antibiotic-induced microbiota depletion disrupts the type 2 immune response critical for biomaterial-mediated tissue repair. Our findings highlight the gut microbiota as a determinant of constructive healing and a potential contributor to inter-individual variability in responses to biologic scaffolds.

## Introduction

The restoration of tissue integrity and homeostasis following injury is a complex and dynamic process influenced by multiple factors. Among these, the immune system is increasingly recognized as a determinant of the quality of tissue repair, orchestrating both the inflammatory response and subsequent regenerative processes (1). In particular, type 2 immunity has emerged as a key driver of tissue regeneration, promoting wound healing, mitigating excessive type 1 inflammation, and facilitating fibroblast and epithelial cell activity (2,3).

Biologically-derived materials, such as extracellular matrix (ECM) scaffolds, have been widely explored for their ability to support a microenvironment conducive to constructive remodeling and tissue repair. These scaffolds enhance biocompatibility and actively regulate cellular function (4). Notably, decellularized ECM has been shown to induce a type 2-mediated immune response, primarily through the production of IL-4, which polarizes macrophages toward an M2 phenotype that promotes tissue repair (5–7). ECM scaffolds, such as porcine urinary bladder matrix (UBM), have been extensively studied in both preclinical and clinical settings for their ability to support tissue regeneration (8–11). These scaffolds provide essential biological cues, immunomodulatory properties, and structural proteins that guide cellular responses and remodeling while also offering mechanical support.

Beyond local tissue factors, systemic influences–including the gut microbiome–play a pivotal role in shaping immune responses during tissue repair. The host-microbiota relationship has co-evolved to maintain immune homeostasis, with symbiotic microbes exerting profound effects on host immunity. The gut microbiota colonizes the gastrointestinal tract contributing to the development of homeostatic mucosal and systemic immune responses, maintenance of epithelial barrier function, and evolution of a balanced microbial ecosystem (12,13). Disruptions of the commensal flora, such as during antibiotic-induced dysbiosis, provide insight into the broader immunological consequences of microbial perturbations. Imbalances in the microbiota-immunity interactions have been linked to extra-intestinal organ dysfunction, including the liver, brain, and lung (14–16). For example, one study found that perinatal administration of streptomycin shifted the gut microbiota toward Bacteroidetes dominance and increased newborn susceptibility to Th1/Th17-driven inflammatory lung disease (17).

The gut microbiota influences immune function, in part, through the production of metabolic by-products derived from bacterial fermentation of dietary components, enzymatic transformations, and interactions with host-derived compounds (18). For instance, one study demonstrated that the antibiotic-induced microbiota depletion (AIMD) model system resulted in a metabolic shift towards anaerobic glycolysis, accompanied by reductions in mitochondrial number and gene expression associated with mitochondrial activity, highlighting the systemic metabolic consequences of microbiome disruption (19). Additionally, microbial metabolites such as butyrate exert anti-inflammatory effects in metabolic-associated steatohepatitis (MASH) by preserving gut barrier function, modulating lipid metabolism, and alleviating insulin resistance (20–22).

Emerging literature suggests that the gut microbiome influences distal tissue healing, fibrotic processes, and immune responses to biomaterials (23–25). Notably, gut-derived RORγ^+^ regulatory T cells, crucial for intestinal homeostasis, were shown to migrate to injured skeletal muscle and regulate repair in a cardiotoxin-induced injury model (25). Another study demonstrated that AIMD slowed the degradation of hyaluronic acid hydrogel implants, reduced immune infiltration, and decreased fibrotic tissue deposition around a silicone implant (23). In a separate study, infection with enterotoxigenic *Bacteroides fragilis* increased neutrophil and γδ T cell infiltration to synthetic polymer polycaprolactone implants, which subsequently altered ECM-related gene expression in fibroblasts during later stages of fibrosis (24). Despite these insights, the specific role of the gut microbiome in modulating type 2 immunity within the context of biological scaffold-driven tissue repair remains largely unexplored.

In this study, we sought to investigate the reciprocal impact of the gut microbiota on the response to MicroMatrix® UBM particulate, a commercially available ECM product used in the clinical setting for tissue repair. Using the AIMD model to disrupt the microbiome and induce dysbiosis, we examined the gut microbiota’s effects on type 2 immune response and tissue remodeling with UBM implantation. Our findings demonstrate that antibiotic treatment reduces type 2 immunity and fosters a pro-fibrotic environment, revealing a critical link between the gut microbiome and the immune-mediated outcomes of ECM-based tissue repair.

## Results

### Treatment with ECM alters the composition and function of the gut microbiota

To investigate the impact of ECM implantation on the composition of the gut microbiome, we coupled a conventionalized germ-free (GF) mouse model with a muscle injury. Germ-free mice, bred in sterile conditions and devoid of any micro-organisms, provide a unique platform to study host-microbe interactions in a controlled and defined context (26). By conventionalizing GF mice with a specific donor microbiota, we can eliminate confounding variables associated with inter-individual microbiome variability and assess the impact of ECM implantation on the microbial community. Briefly, we conventionalized C57BL/6 GF mice with a young murine fecal microbiota transplant (FMT) and allowed two weeks for colonization. After colonization, we performed bilateral volumetric muscle loss (VML) injuries in the quadriceps and implanted UBM, a decellularized ECM scaffold, in the defect space. Control groups included naïve mice (no injury) and VML-injured mice receiving saline. Stool samples were collected at days 0, 7, 21, and 42 post injury for 16S rRNA V3-V4 gene sequencing (Fig. 1A).

**Figure 1.**
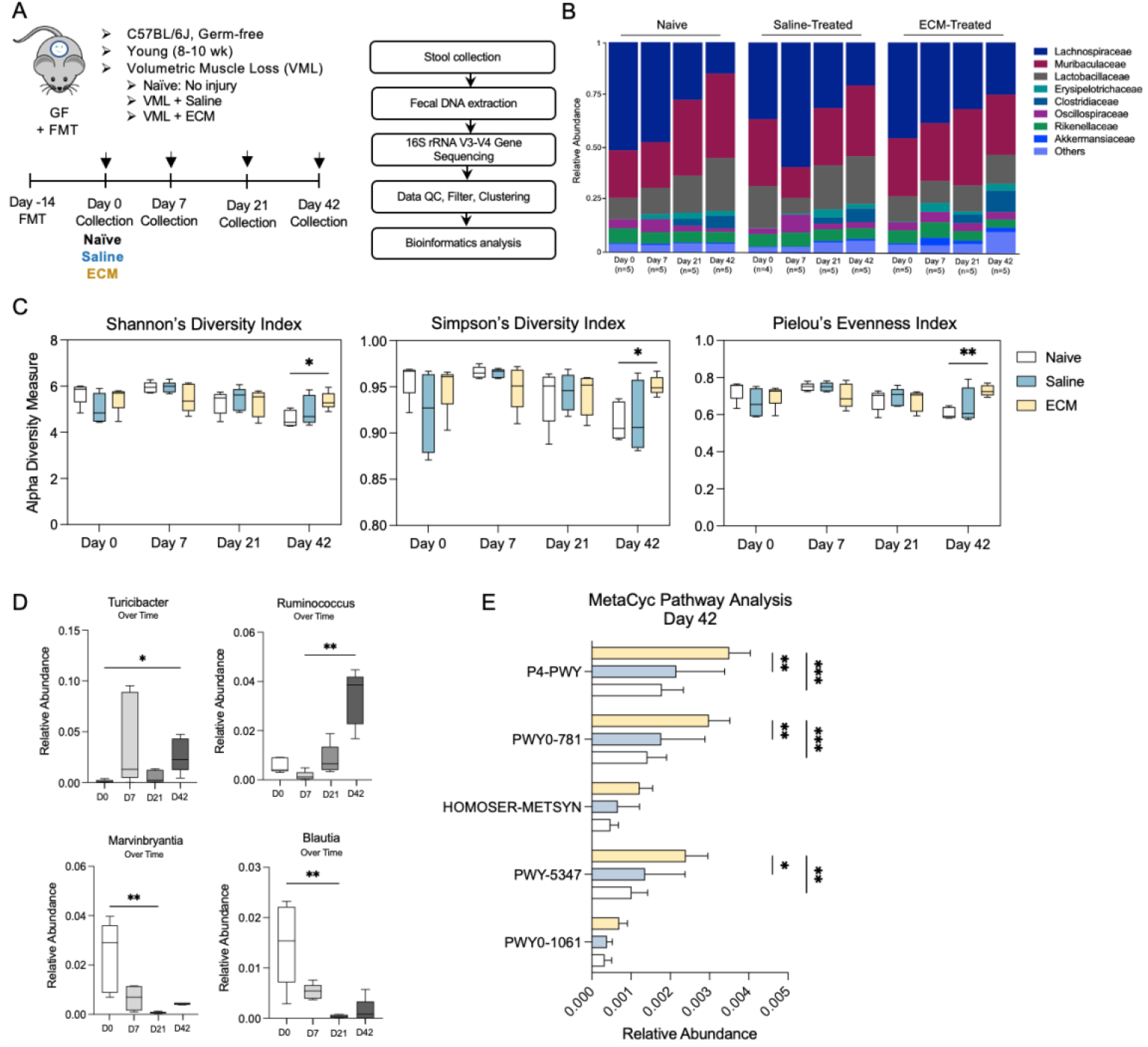
ECM treatment alters the composition and function of the gut microbiome. (A) Schematic of experimental timeline consisting of no injury naïve control, volumetric muscle loss (VML) injury with saline, and VML treated with extracellular matrix (ECM) in conventionalized germ-free mice (GF+FMT). Arrows depict stool collection timepoints for 16S rRNA V3-V4 gene sequencing. (B) Family-level microbiota composition of naïve, saline, and ECM-treated mice over time expressed as relative abundances. (C) α-Diversity of the gut microbiome in naïve, saline, and ECM-treated mice as measured by Shannon’s and Simpson’s indices and Pielou’s Evenness index. (D) Quantification of the relative abundance of select genus-level bacteria in ECM-treated mice over time. (E) Quantification of the relative abundance of selected MetaCyc pathways from PICRUSt2 analysis in naïve, saline, and ECM-treated mice at day 42-post VML-ECM. (D) Kruskal-Wallis test with Dunn’s multiple comparisons test. (C, E) Ordinary two-way ANOVA with Tukey’s multiple comparisons test. *P < 0.05, **P < 0.01, ***P < 0.001.

Using 16S rRNA sequencing, we characterized the temporal dynamics of microbial communities following conventionalization. Plotting relative abundance of bacterial families over time revealed an evolving population in naïve mice. Lachnospiraceae and Muribaculaceae were the most abundant family of bacteria in the conventionalized mice (Fig. 1B), consistent with other studies on laboratory bred mice (27). To further characterize microbial community structure, we assessed the bacterial richness and evenness in the naïve, saline-treated, and ECM-treated groups across all time points. Analysis of alpha diversity using Shannon’s and Simpson’s diversity index revealed a significant increase in microbial diversity in the ECM-treated group compared to naïve mice at day 42 (Fig. 1C). Pielou’s Evenness index demonstrated a significant increase in species evenness in ECM-treated mice at the day 42 time point compared to naïve controls, indicating a more equitable distribution of bacterial abundances within the community (Fig. 1C). There were no changes in the Observed and Chao1 alpha diversity measures (SI Appendix, Fig. S1A). These findings suggest that injury and ECM implantation may contribute to long-term shifts in microbiome diversity, evenness, and community structure.

Building on our observations of altered microbial diversity following ECM implantation, we next examined taxonomic shifts at the genus level to identify specific microbial populations associated with ECM treatment. Linear discriminant analysis (LDA) revealed that at day 7 post-ECM implantation, *Turicibacter* were most abundant, while at day 42 post-ECM implantation *Ruminococcus* expanded (SI Appendix, Fig. S1B). Specifically, *Turicibacter* and *Ruminococcus* were significantly enriched in ECM-treated groups at the day 42 time point compared to day 0 baseline (Fig. 1D). These genera were also more abundant in ECM-treated mice compared to naïve and saline-treated groups at day 42, albeit non-significantly (SI Appendix, Fig. S1C). Additionally, we observed a decrease in *Marvinbryantia* and *Blautia* in ECM-treated groups over time (Fig. 1D). Notably, prior studies have reported increased *Marvinbrynatia* and *Blautia* abundance in mice implanted with silicone, suggesting that these genera may be associated with implant-specific responses (23). However, previous studies have also shown that *Ruminococcus* decreased in response to silicone implantation (23), opposite to the trend we observed with ECM treatment.

To investigate bacterial signaling pathways associated with compositional alterations, we utilized PICRUSt2 to predict MetaCyc metabolic pathways. We identified enrichment in the relative abundance of bacteria that participate in pathways associated with the biosynthesis of amino acids, L-phenylalanine (P4-PWY), L-arginine (PWY0-781), L-ornithine (PWY0-781), L-methionine (HOMOSER-METSYN-PWY), L-isoleucine (PWY-5347), L-valine (PWY-5347), and L-leucine (PWY-5347) in the ECM-treated condition at day 42 compared to naive (Fig. 1E). Amino acids, like phenylalanine, are essential for epithelial cell repair within the gut and regulating intestinal immunity (28,29). It has been reported that P4-PWY, PWY0-781, and HOMOSER-METSYN-PWY are decreased in inflammatory gut conditions including inflammatory bowel disease (IBD) and colorectal cancer (CRC) (30). These findings suggest that ECM implantation may influence the utilization of host amino acids by intestinal bacteria.

### Antibiotic treatment decreases pro-regenerative macrophages and IL-4-producing CD4^+^ T cells in the ECM implant microenvironment

To determine whether the gut microbiota influences the type 2 regenerative immune response to a potent type 2-driven biological scaffold, we utilized the AIMD model system. This system has been used to disrupt the composition of the microbiome, reduce diversity, and alter metabolism (31,32). It has also been employed to elucidate the microbiome’s role in pathological disease, particularly across diverse injury models (33–36). To assess the impact of the gut microbiota on the type 2 response of ECM implants, we induced dysbiosis by administering four broad-spectrum antibiotics–ampicillin, neomycin, vancomycin, and metronidazole –via drinking water. We performed bilateral VML surgeries in the quadriceps and implanted an ECM scaffold in the defect space. To integrate AIMD with the VML system, we initiated a 7-day antibiotic regimen beginning on the day of surgery (Fig. 2A). Tissue was collected and analyzed at either day 7-post VML-ECM to characterize immune dynamics during the peak inflammation or day 42-post VML-ECM to evaluate tissue remodeling and healing.

**Figure 2.**
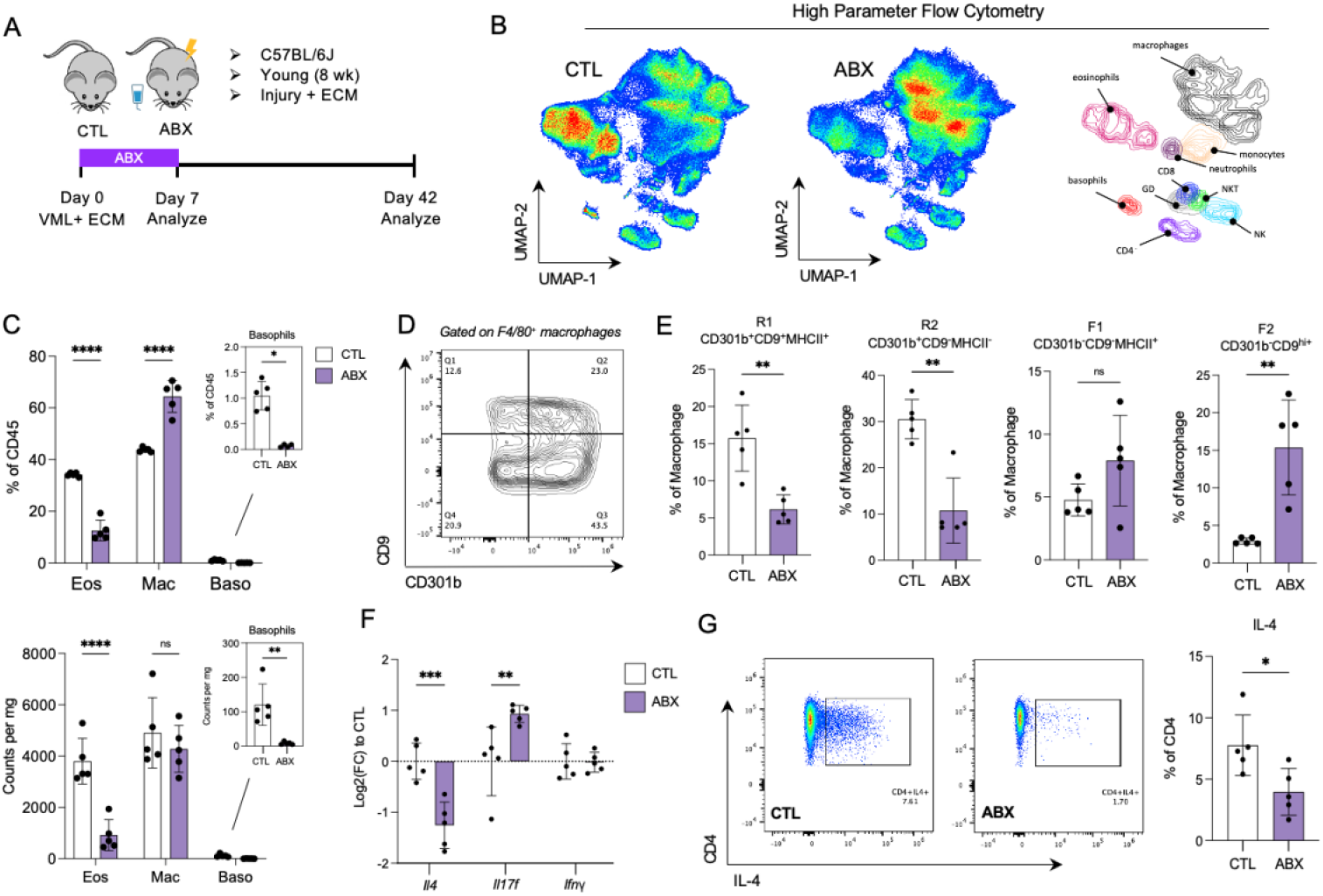
Depletion of the gut microbiota alters type 2 immunity to an ECM scaffold. (A) Schematic of experimental timeline coupling antibiotic-induced microbiota depletion and volumetric muscle loss (VML) surgery treated with extracellular matrix (ECM) scaffold. Abx = antibiotics (vancomycin, neomycin, metronidazole, and ampicillin) (see Methods for details). (B) Overall immune infiltrate of ECM-treated muscle at day 7-post VML-ECM detected by flow cytometry and displayed by dimensionality reduction algorithms, Uniform Manifold Approximation and Project (UMAP). (C) Flow cytometric quantification of Eos=Eosinophils, Mac=Macrophages, and Baso=Basophils of ECM-treated muscle at day 7-post VML-ECM. (D) Representative gating scheme of R1 (CD301b^+^, CD9^+^, MHCII^+^), R2 (CD301b^+^, CD9^-^, MHCII^-^), F1 (CD301b^-^, CD9^-^, MHCII^+^), and F2 (CD301b^-^, CD9^hi+^) macrophages (see SI Appendix, Fig. S8 for details). (E) Flow cytometric quantification of R1, R2, F1, and F2 macrophages of ECM-treated muscle at day 7-post VML-ECM. (F) Quantification of *Il4, Il17f*, and *Ifny* gene expression using qRT-PCR in ECM-treated muscle at day 7-post VML-ECM. (G) Representative flow cytometric images (left) and quantification (right) of IL-4-producing CD4^+^ T cells in ECM-treated muscle at day 7-post VML-ECM. Bar graphs show mean ± SD. Data are representative of *n* = 5 mice. (C, F) Two-way ANOVA with Sidak’s multiple comparisons. (E, G) Mann-Whitney test. ns=not significant; *P < 0.05, **P < 0.01, ***P < 0.001.

To evaluate broad immunological changes at day 7-post VML, we performed high parameter flow cytometry (SI Appendix, Fig. S8) on the VML-injured, ECM-implanted quadricep muscle (Fig. 2B). We observed a significant decrease in CD45^+^ immune cells in the antibiotic-treated group, both as a percentage of live cells and as counts per milligram of tissue (SI Appendix, Fig. S2A). The administration of antibiotics resulted in a significant decrease in eosinophils (CD45^+^CD11b^+^SiglecF^+^MHCII^-^) and basophils (CD45^+^CD11b^+^CD200R3^+^) (Fig. 2C). Macrophages (CD45^+^CD11b^+^Ly6CmidF4/80^+^) were significantly increased as a percentage of CD45^+^ immune cells, however, this was not reflected in the counts (Fig. 2C). We further profiled these macrophages based on previous work that has identified regenerative (R1 and R2) and fibrotic (F1 and F2) macrophages in the context of biomaterial implantation (37). Based on single cell analysis, R1 macrophages are functionally characterized by glycolysis, whereas R2 macrophages rely on type 2-like oxidative phosphorylation (37). F1 macrophages are characterized by type I inflammation, while F2 are associated with type 17 inflammation (37). Pro-regenerative macrophages are characterized by the expression of surface markers CD301b, CD9, and MHCII (Fig. 2D). Interestingly, the frequency of pro-regenerative R1 (CD301b^+^CD9^+^MHCII^+^) and R2 macrophages (CD301b^+^CD9^-^MHCII^-^) were significantly decreased in the antibiotic-treated groups (Fig. 2E). This reduction was reflected in the counts as well (SI Appendix, Fig. S2C). Furthermore, the frequency of fibrotic macrophage populations, F1 (CD301b^-^CD9^-^MHCII^+^) and F2 (CD301b^-^ CD9^hi+^) were increased with antibiotics (Fig. 2E).

To investigate the polarization of T cells in the ECM-treated quadricep, we first looked at the gene expression of *Ifnγ, Il4*, and *Il17f* in the bulk quadricep to assess canonical TH1, TH2, and T*H*17 activity, respectively. We observed a significant decrease in *Il4* and subsequent increase in I*l17f* gene expression with antibiotic treatment at day 7-post VML-ECM (Fig. 2F). To explore the source of these cytokines further, we performed intracellular cytokine staining on the VML-injured, ECM implanted quadricep muscle, and found that the frequency of IL-4-producing CD4^+^ T cells were decreased in the antibiotic group (Fig. 2G). We did not observe any changes in IL-17A-producing CD4^+^ T cells in the quadriceps; however, IL-17A-producing CD4^+^ and γδ T cells showed an increasing trend in the inguinal lymph nodes (SI Appendix, Fig. S3D).

Taken altogether, these findings demonstrate that antibiotic-induced microbiota depletion impacts the local type 2 immune response in the context of UBM implantation at the height of the inflammatory response (day 7) following VML-ECM. This suggests that the dysbiosis of the gut microbiota influences the distal immunological microenvironment of a biological scaffold by shaping both immune cell composition and activation.

### Antibiotic treatment disrupts the composition of the gut microbiome and impacts the colon microenvironment

Next we sought to understand how AIMD with ECM implantation impacts the gut microenvironment. To assess microbiome dynamics, we performed 16S rRNA sequencing on stool samples collected from control and antibiotic-treated mice with ECM implants at multiple time points (day 0-, 7-, 21-, and 42-post antibiotic treatment). At the genus-level, the microbiome composition was significantly disrupted by day 7, with a notable outgrowth of *Enterococcus* and *Staphylococcus* (Fig. 3A). Principle co-ordinates analysis (PCoA) revealed that the microbial community composition at day 7-post antibiotic treatment was distinct from other time points, with this altered community structure persisting through day 21 and 42 (Fig. 3B, SI Appendix Fig. S4A). Furthermore, Shannon’s diversity index indicated a significant reduction in alpha diversity at day 7, which remained consistently lower at day 21 and 42, though not statistically significant at later time points (Fig. 3C). Consistent with this, both Simpson’s and Pielou’s evenness indices showed significant reductions at day 7 (SI Appendix, Fig. S4B).

**Figure 3.**
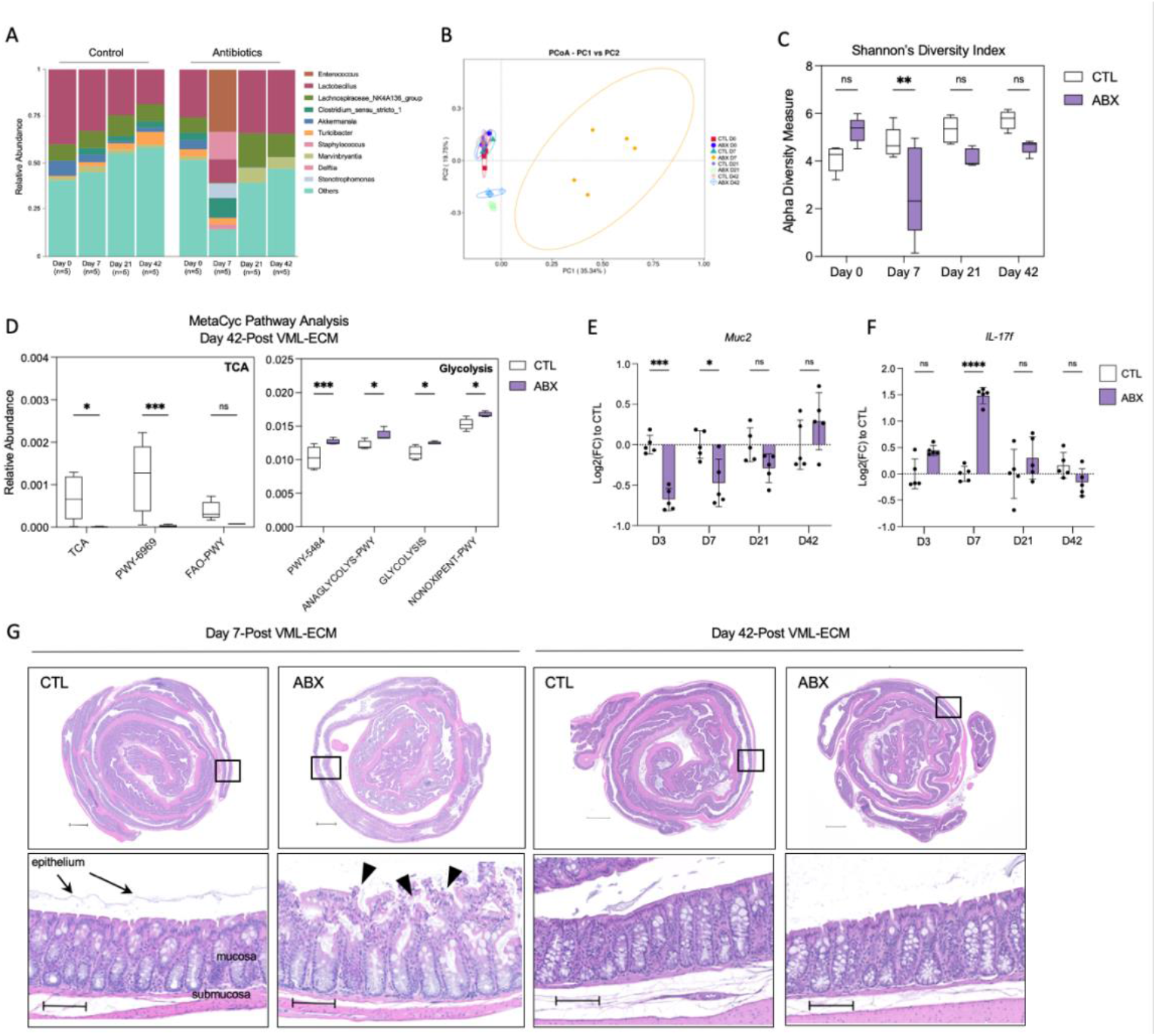
Antibiotic treatment disrupts the composition of the gut microbiome and epithelial integrity. (A) Genus-level microbiota composition of control and antibiotics-treated mice with VML-ECM over time expressed as relative abundances. (B) Principle co-ordinates analysis (PCoA) of unweighted and normalized UniFrac distances of microbiota from control and antibiotics-treated mice with VML-ECM at several time points. (C) α-Diversity of the gut microbiome in control and antibiotic-treated mice with VML-ECM at several time points as measured by Shannon’s index. (D) Quantification of the relative abundance of bacteria that participate in selected MetaCyc pathways from PICRUSt2 analysis in control and antibiotics-treated mice at day 42 post VML-ECM. (E) Quantification of *Muc2* and *Il17f* gene expression using qRT-PCR in control and antibiotics-treated colon tissue over time. (F) H&E image of the colon at day 7- and 42-post VML-ECM in control and antibiotic-treated mice. Top row scale bar = 1 mm. Bottom row scale bar = 100 μm. Bar graphs show mean ± SD. Data are representative of *n* = 5 mice. (C, D, E) Ordinary two-way ANOVA with Sidak’s multiple comparisons test. *P < 0.05, **P < 0.01, ***P < 0.001.

To further characterize functional alterations within the microbiome, we utilized PICRUSt2 analysis for MetaCyc pathways at day 42-post antibiotic treatment. We observed a significant reduction in the abundance of bacteria that contribute to pathways associated with tricarboxylic cycle (TCA), including TCA Cycle I (TCA), TCA Cycle V (PWY-6969), and fatty acid β-oxidation I (FAO-PWY), in the antibiotic treated group (Fig. 3D). Conversely, there was a significant upregulation of glycolysis-associated pathways, including glycolysis II (PWY-5484), glycolysis III (ANAGLYCOLYS-PWY), glycolysis I (GLYCOLYSIS), and pentose phosphate pathway (NONOXIPENT-PWY), suggesting a metabolic shift favoring anaerobic glycolysis over oxidative phosphorylation (Fig. 3D).

To assess local changes in the colonic microenvironment, we performed qRT-PCR on bulk colon tissue at days 3, 7, 21, and 42, focusing on genes associated with barrier health and function. We detected a significant decrease in *Muc2* at days 3- and day 7-post antibiotic administration (Fig. 3E). *Muc2* encodes a critical mucin produced by goblet cells that maintains the mucus layer and protects against inflammatory insults (38). Additionally, *Reg3γ* expression was significantly reduced at day 3-post antibiotics (SI Appendix, Fig. S4C). REG3γ is an antimicrobial peptide that selectively targets gram-positive bacteria and plays a role in limiting bacterial translocation across the intestinal barrier (39). In contrast, we observed a significant increase in *Il17f* gene expression at day 7, suggesting a transient pro-inflammatory response (Fig. 3F). These alterations in barrier function and immune activation gradually resolved by days 21 and 42, returning to near-homeostatic levels. Histological examination of Swiss-rolled colonic sections stained with hematoxylin and eosin (H&E) revealed substantial epithelial barrier disruption at day 7, supporting the observed molecular changes seen with qRT-PCR. By day 42, epithelial integrity, as assessed by H&E staining, improved in the antibiotic-treated group, mirroring the transcriptional recovery of barrier-associated genes (Fig. 3G).

### Antibiotics reduce myocytes with centrally located nuclei and promote pro-fibrotic transcriptional activity at the ECM implant site

Centrally located nuclei are commonly used to assess regenerating myofibers. Under homeostatic conditions, skeletal muscle is defined by the hallmark peripheral positioning of myonuclei. In contrast, regenerating muscle fibers exhibit centrally located nuclei due to the fusion of activated satellite cells during muscle repair, initially positioning the nuclei centrally before they migrate to the periphery as the muscle matures (40). We performed dystrophin immunofluorescence staining to visualize individual myocytes and DAPI staining to quantify centrally located nuclei at day 42-post VML-ECM. We observed a reduction in centrally located nuclei as a percentage of total myocytes in antibiotic-treated conditions suggesting a reduced capacity to regenerate (Fig. 4A).

**Figure 4.**
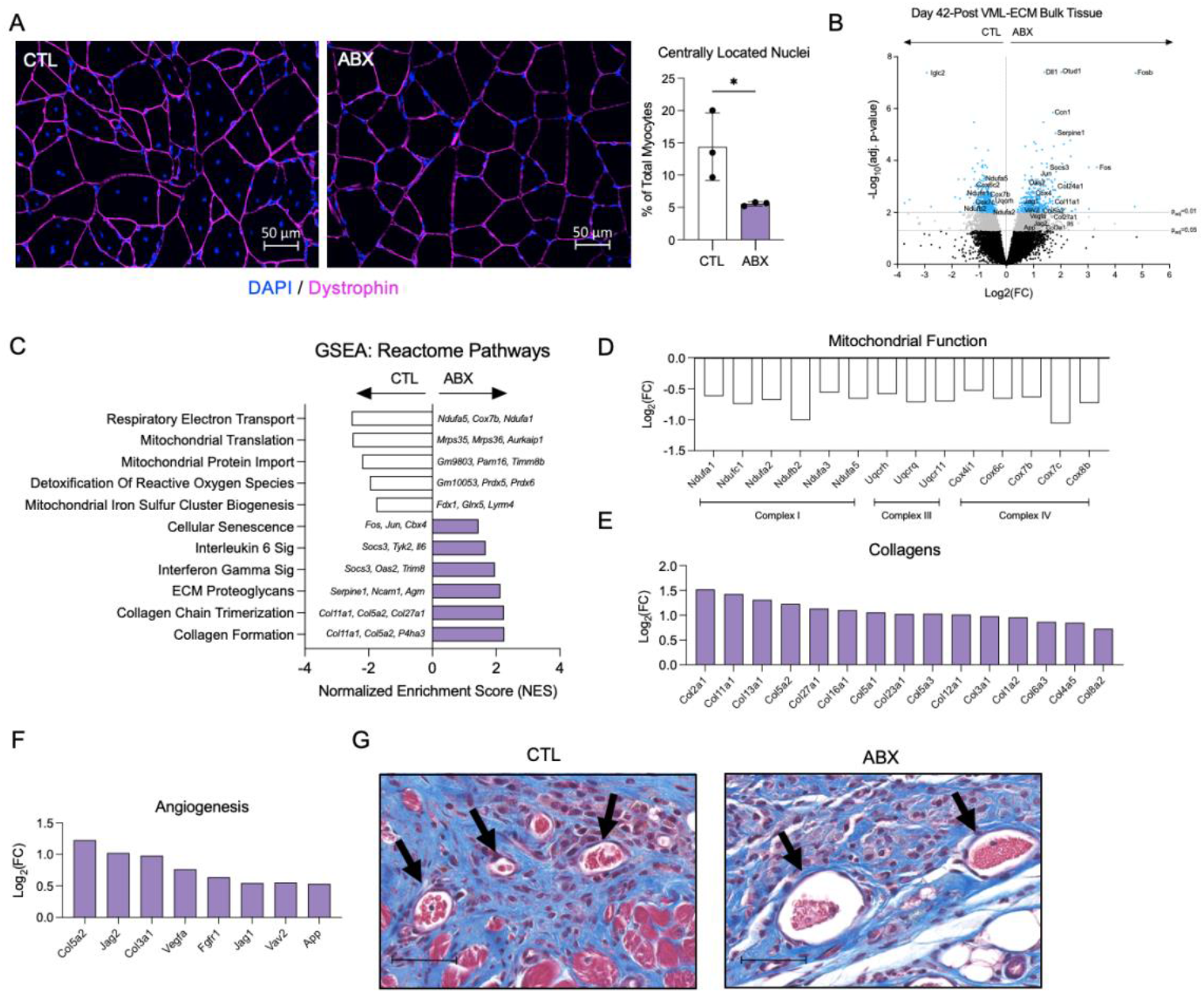
Antibiotics impair the pro-regenerative response of an ECM scaffold. (A) Immunofluorescence staining (left) of DAPI (blue) and Dystrophin (purple) of ECM-treated quadricep tissue (axial cut) in control and antibiotic treated mice at day 42-post VML-ECM. Scale bar = 50 μm. Quantification (right) of centrally located nuclei based on DAPI and dystrophin immunofluorescence staining. (B) Volcano plot of differential gene expression results of bulk RNA sequencing from ECM implant site at day 42-post VML-ECM. (C) Quantification of selected Reactome pathways from gene set enrichment analysis (GSEA) with normalized enrichment scores (NES). (D) Manual categorization of significant differentially expressed genes from bulk sequenced ECM implant at day 42-post VML+ECM related to mitochondrial function. (E) Manual categorization of significant differentially expressed collagens from bulk sequenced ECM implant at day 42-post VML-ECM. (F) Manual categorization of significant differentially expressed genes of bulk sequenced ECM implant at day 42-post VML-ECM related to angiogenesis. (G) Representative Masson’s Trichrome staining of ECM-treated injured quadricep tissue (axial cut). Scale bar = 50 μm. (A) Unpaired t-test. Bar graph shows mean ± SD. *P < 0.05, **P < 0.01, ***P < 0.001.

Due to this reduction in regenerating myocytes, we performed bulk RNA-sequencing on ECM-implanted quadricep tissue to better understand the transcriptional activity within the tissue site. Bulk RNA sequencing of the injured quadricep implanted with ECM resulted in 3,346 significantly differentially expressed genes (DEGs). Of those, 1,783 genes were upregulated, and 1,563 genes were downregulated with antibiotic treatment (Fig. 4B). The most significantly upregulated gene were *Dll1* and *Fosb*, while the most significantly downregulated was *Iglc2. Dll1*, a homolog of notch delta ligand, is part of the intracellular Notch signaling pathway. While the careful regulation of Notch pathways participates in myocyte regeneration and stem cell differentiation, its aberrant upregulation has also been associated with inflammatory signaling that promotes chronic fibroblast activation and is recently a target for anti-fibrotic therapy (41,42). Downstream of such inflammatory cascades, *Fosb* encodes for a transcription factor that dimerizes with Jun to form the AP-1 complex. The chronic overexpression of the AP-1 has also been implicated in excessive collagen production and pathological fibrosis (43,44).

Gene set enrichment analysis (GSEA) of the Reactome and Hallmark pathways revealed several significant pathways related to tissue repair. Notably, the antibiotic-treated group exhibited a significant increase in pathways for ECM Proteoglycans, Collagen Chain Trimerization, and Collagen Formation, suggesting enhanced fibrosis, a process commonly characterized by excessive collagen deposition (Fig. 4C). Furthermore, Interferon Gamma signaling and Interleukin 6 signaling were upregulated with antibiotics (Fig. 4C). Pathways associated with mitochondrial function, including respiratory electron transport, mitochondrial translation, mitochondrial protein import, and the detoxification of reactive oxygen species were all downregulated with antibiotic treatment (Fig. 4C). Interestingly, GSEA of the Hallmark pathways revealed downregulation of oxidative phosphorylation and Mtorc1 signaling with antibiotics treatment (SI Appendix, Fig. S5A). To further investigate mitochondrial dysfunction, we manually categorized genes associated with complex I (*Ndufa1, Ndufc1, Ndufa2, Ndufb2, Ndufa3*, and *Ndufa5*), complex III (*Uqcrh, Uqcrq*, and *Uqcr11*), and complex IV (*Cox4i1, Cox6c, Cox7b, Cox7c*, and *Cox8b*). We observed a notable downregulation of these genes with antibiotics treatment, suggesting impaired mitochondrial function (Fig. 4D). In contrast, manual categorization of collagen-related genes revealed several significantly upregulated collagens, including *Col2a1, Col11a1, Col13a1, Col5a2*, and *Col27a1* (Fig. 4E). After observing the changes in gene expression of collagen genes, we performed Masson’s Trichrome and Picrosirius Red staining on quadriceps implanted with ECM and quantified fibrosis within the quadricep. There were no significant differences in fibrosis between the control and antibiotic-treated groups at day 42-post VML-ECM (SI Appendix, Fig. S6A-D).

The formation of new blood vessels during tissue repair is a vital process to constructive tissue remodeling by supplying essential nutrients, immune cell access, and oxygen. However, the precise timing and regulation of the angiogenic response, particularly during the transition from inflammation to remodeling, are key factors for ensuring effective repair and preventing pathological outcomes (45). Recognizing its significance, we investigated the expression of several angiogenesis-associated genes and observed significant upregulation of *Col5a2, Jag2, Col3a1, Vav2, Vegfa, Fgfr1*, and *App* (Fig. 4F). Based on these differentially expressed genes, we performed Masson’s Trichrome staining to observe the architecture of blood vessels within the interface of the ECM implant and quadricep. Notably, antibiotic-treated mice exhibited enlarged vessel size compared to the control group (Fig. 4G).

## Discussion

Tissue repair mediated by biological scaffolds is intricately linked to the immune system. Following injury, a sequence of events triggers the acute inflammatory phase, characterized by the migration of immune cells from systemic circulation to begin the clearance of cellular debris and initiate vascularization and tissue deposition. In the following weeks, the response shifts toward chronic inflammation and tissue repair, marked by macrophage polarization and T cell activation (46,47). If left unresolved, this process can lead to excessive collagen deposition and fibrosis. The gut microbiome is increasingly recognized as a key modulator of the immune response, shaping systemic immunity by inducing T helper 17 and T regulatory cells, influencing the host antibody repertoire, and regulating the production of several innate immune cells in distal organs (48–54). Emerging evidence suggests that the gut microbiome impacts tissue repair across multiple organs, with studies demonstrating its role in modulating wound healing, muscle regeneration, and fibrotic processes (23,25).

The findings presented here contribute to this growing body of evidence by demonstrating that implantation of ECM alters gut microbiota composition and metabolic function, while also influencing distal immune response and tissue repair outcomes. Notably, we show that ECM implantation leads to increased microbial diversity, enrichment of specific bacterial genera, and functional shifts in metabolic pathways related to amino acid biosynthesis (Fig. 1C-E). These microbial changes correspond to immune and regenerative responses, highlighting the interplay between the microbiome and host tissue repair.

Using a model of AIMD, we demonstrate that disruption of the gut microbiome significantly impacts type 2 immunity and impairs the pro-regenerative response to an ECM scaffold. Specifically, AIMD resulted in a decrease in pro-regenerative macrophage populations (R1 and R2) and IL-4-producing CD4^+^ T cells, while promoting the expansion of fibrotic macrophages (F1 and F2) and increasing *Il17f* expression (Fig. 2E-G). The IL17 family plays a prominent pro-inflammatory role in gut and connective tissues. Though IL17 contributes to host defense and acute tissue repair, its sustained elevation has been associated with a pathological role in chronic healing and development of fibrosis (55). Notably, *Il17f* expression peaked in the colon at day 7-post AIMD, suggesting a temporally regulated immune response that may contribute to persistent inflammation and impaired healing at the distal injury site. Furthermore, we observed a significant increase in the pro-fibrotic macrophage, F2, whose function has been previously shown to be associated with type 17 inflammation (37). Given the established role of IL17 signaling in aberrant healing processes, future studies should further investigate its mechanistic contributions to the impaired response to ECM in the absence of a balanced microbiome. These findings underscore the critical role of the microbiome in shaping distal immune responses to biomaterials and suggest that an intact microbiome supports a regenerative tissue microenvironment.

Our bulk sequencing analysis at day 42-post VML-ECM revealed changes in several pathways and genes associated with constructive tissue repair. We saw a significant reduction in pathways and genes associated with mitochondrial function, particularly with respiratory electron transport and oxidative phosphorylation (Fig. 4C-D). Glycolysis and oxidative phosphorylation are tightly linked metabolic pathways that dynamically shift during tissue repair, with glycolysis typically dominating in early inflammatory phases before transitioning to oxidative phosphorylation (56,57). Interestingly, a regenerative macrophage, R2, was significantly decreased at the day 7 timepoint (Fig. 2E), and has been previously shown to possess oxidative phosphorylation function based on single cell characterization (37). It has been shown that antibiotics, particularly bactericidal antibiotics, can cause mitochondrial dysfunction in mammalian cells resulting in elevated oxidative stress and reactive oxygen species (ROS) overproduction (58). Further studies are needed to determine the functional consequences of impaired oxidative phosphorylation in the context of ECM-implantation and whether metabolic interventions can restore a pro-regenerative environment. Future investigations could include metabolic flux analysis to assess real-time shift in cellular energy production and *in vivo* studies testing pharmacological or genetic approaches to enhance mitochondrial function.

The gut microbiota exerts a profound influence on systemic immunity through several distinct mechanisms. First, microbial molecules–such as microbially-derived metabolites or structural components of bacterial cell walls–can enter systemic circulation and directly engage immune cells. For example, a microbially-derived peptidoglycan has been shown to activate an innate pattern recognition receptor, Nod1, to activate neutrophils and influence their cytotoxic function (59). Second, the microbiota continuously primes the immune system for cytokine production, which can exert widespread systemic effects. One study demonstrated that the commensal microbiota instructs plasmacytoid dendritic cells to produce type I interferons utilizing germ-free and monocolonized germ-free mice (60). Lastly, immune cells educated within the gut can migrate to peripheral tissues, extending its influence to extraintestinal organs (48–50). In the AIMD model, we observed that antibiotic-treated mice exhibited a significant decrease in the CD4:CD8 T cell ratio in the blood of antibiotic-treated mice (SI Appendix, Fig. S7A) and a significant increase in CD3^+^ T cells in the spleen on day 7 (SI Appendix, Fig. S7B). These findings suggest that antibiotics alter systemic T cell composition, potentially contributing to impaired type 2 function at the ECM-VML site. Understanding the mechanistic link between the ECM-treated VML site and the gut microbiome can better inform the advancement of immunomodulatory strategies. Monocolonization studies using candidate bacteria found in this study could help identify species that enhance or impair tissue repair. Furthermore, given prior evidence that aging reduces the regenerative capacity of ECM (61), our findings suggest that alterations in the aging microbiome may contribute to this decline. The potential impact of age-related microbiome changes on ECM-mediated regeneration warrants further investigation. Harnessing this connection could lead to microbiome-targeted therapies that optimize immune responses for improved tissue repair and regeneration.

Despite the valuable insights gained from this study, several limitations inherent to microbiome research should be considered. Microbiome composition varies significantly across different animal facilities, and the origin of conventionalized microbiota can result in cage effect that influences experimental outcomes (62). In our study, naïve mice exhibited dynamic shifts in bacterial populations over time (Fig. 1B), highlighting the challenges of standardizing microbiome studies and comparing phylogenetic changes across different studies. Additionally, the AIMD model has its own limitations, including the duration and timing of antibiotic administration and variations in antibiotic consumption among individual mice (63). While antibiotics have revolutionized infectious disease treatment, they also induce long-term disruptions in the gut microbiome that may persist for decades. The consequences of antibiotic-induced dysbiosis on human health remain an active area of investigation. It is important to note that AIMD does not have a complete depletion of the gut microbiome and antibiotics themselves can have a direct effect on host cells (64). Future studies should examine the effects of individual antibiotics from the AIMD model system on type 2 immunity and tissue repair to determine whether specific antibiotics drive particular immunological and regenerative outcomes. Alternatively, the use of germ-free mice may provide additional support for the findings presented here; however, these mice exhibit developmental immune deficiencies that may complicate interpretation (26). The strength of the AIMD model lies in its use of mice with preserved immune development and function. Additionally, antibiotics were administered at the time of injury with implantation to better reflect clinical scenarios.

In conclusion, our study highlights the critical role of the gut microbiome in modulating immune responses and tissue regeneration following ECM implantation. By demonstrating that microbiota depletion shifts immune cell composition, disrupts metabolic pathways, and alters tissue repair outcomes, we provide compelling evidence for the microbiome’s influence on biomaterial-mediated healing. Future studies should explore microbiome-targeted interventions as potential strategies to enhance regenerative medicine therapies and optimize patient outcomes.

## Materials and Methods

### Mice

All animal procedures were approved by Johns Hopkins University Institutional Animal Care and Use Committee protocol. Mice were housed and maintained in the Johns Hopkins Cancer Research Building animal facility for antibiotic studies or the Johns Hopkins Germ-Free Mouse Core for conventionalized germ-free studies. Seven-to eight-week-old female C57BL/6 were obtained from Jackson Laboratory (Strain #00064) were utilized for all antibiotic-induced microbiota depletion experiments. Seven-to ten-week-old female C57BL/6 were obtained from the Germ-Free Mouse Core for all conventionalizing germ-free experiments.

### Conventionalized germ-free model using fecal microbiota transplantation (FMT)

Three groups of five female GF C57BL/6 mice were colonized by oral gavage (200 μl/mouse) with murine gut microbiota. Murine stool samples were acquired from seven-ten-week old C57BL/6 mice from the National Institute on Aging and snap frozen on dry ice and subsequently frozen at -80°C until use. GF mice received a fecal microbiota transplant (FMT) of a 4% w/v stool slurry in sterile 1X Dulbecco’s phosphate buffered saline (DPBS) (Gibco) by oral gavage. Mice were conventionalized with the FMT for two weeks prior to starting surgical experiments. Fecal samples for 16S rRNA amplicon sequencing were collected by individual clean catch at the start of the experiment before receiving volumetric muscle loss surgery, then subsequently at 7-days, 21-days, and 42-days-post surgery. Stool samples were snap frozen on dry ice and subsequently frozen at -80°C until DNA extraction.

### Antibiotic-induced microbiota depletion (AIMD) model

C57BL/6 female mice were randomized into two groups: control or antibiotics. For the induction of antibiotic-induced microbiota depletion, mice were treated with broad-spectrum antibiotics in drinking water. Mice were treated with antibiotics for one week. The antibiotics cocktail was comprised of four antibiotics: ampicillin (1g/L; MWI), vancomycin (500 mg/L), metronidazole (1g/L; MWI), and neomycin (1g/L) dissolved in sterile distilled water provided ad libitum via regular drinking bottles. Control animals were given sterile distilled water.

### Stool collection, DNA isolation, and 16S rRNA amplicon sequencing

Fecal samples were collected by individual catch clean, snap frozen on dry ice, and subsequently frozen at -80°C until use. DNA for 16S rRNA amplicon gene sequencing was isolated using Quick-DNA Fecal/Soil Microbe Miniprep Kit (Zymo). The DNA isolation steps were performed according to the manufacturer protocol. Amplicon library building of the V3-V4 hypervariable region, data quality control, and sequencing was performed by Novogene.

### Volumetric muscle loss (VML) model and biological scaffold preparation

Bilateral muscle defects in the murine quadricep were created as previously described (5). Briefly, mice were anesthetized and maintained using 2-2.5% isoflurane. 5 mg/kg of carprofen (Rimadyl, MWI Animal Health) was injected subcutaneously for pain relief. Hair was removed above the quadricep with an electric razor (Oster) and the area was sterilized with 70% ethanol. After a 1-1.5-cm skin incision was created above the quadricep muscle, approximately 3 mm x 4 mm x 4 mm segment of quadricep was resected using surgical scissors to create a defect space. The bilateral defects were filled with 50 μl/quadricep of 200mg/mL of MicroMatrix® Urinary Bladder Matrix (Integra) hydrolyzed in sterile 1X Dulbecco’s phosphate buffered saline (DPBS) (Gibco) acquired from Integra. Control surgeries for the conventionalized GF mouse study were treated with 50μl/quadricep of sterile DPBS. The skin incision was closed through the use of surgical sutures. Mice were euthanized at day 7- or 42-post surgery for analysis.

### qRT-PCR

#### Tissue preparation, RNA extraction, cDNA synthesis, and qPCR

Harvested tissue were immediately placed into TRIzol (Invitrogen) after mouse euthanasia. All samples were subsequently snap frozen on dry ice and stored at -80°C until use. Prior to starting RNA extraction, samples were thawed on ice and homogenized with approximately six to eight ceramic beads (2.8 mm; OMNI International) using the Bead Ruptor 12 (OMNI International). The Bead Ruptor was programmed to the highest speed for 3 rounds of 15 seconds. Subsequent chloroform extraction was utilized on the homogenized samples. The RNeasy PLUS Mini Kit (Qiagen) was used for RNA isolation and purification according to manufacturer’s instructions. Quantification of RNA was performed using the NanoDrop 2000 (ThermoFisher Scientific). cDNA synthesis occurred using SuperScript IV VILO Master Mix (ThermoFisher Scientific) and a C100 Touch Thermocycler (BioRad). qRT-PCR was performed on the StepOne Plus Real-Time PCR System and software (Applied Biosystems, ThermoFisher Scientific) using TaqMan Gene Expression Master Mix (Applied Biosystems) and TaqMan probes (listed in SI Appendix, Table S1) according to manufacturer’s instructions. For quadricep tissue samples, *Rer1* was used as an endogenous control (reference housekeeping gene). For colon tissue samples, *Gapdh* was used as an endogenous control (reference housekeeping gene). Samples were normalized to control (no antibiotic treatment) unless otherwise stated. All data was analyzed using the 2^-ΔΔCt^ Ct method (65).

### Flow cytometry

#### Tissue preparation

Quadricep muscle with UBM implantation were harvested by cutting the quadricep from the hip to the knee. The tissue was finely diced and digested for 45 minutes at 37°C with 1.67 Wunsch U/mL (5mg/mL) Liberase TL (Roche Diagnostics) and DNase I (0.2 mg/mL) (Gibco) in RPMI 1640 medium with L-Glutamine (Gibco) and 25 mM HEPES (Gibco). Digested tissue was grinded through a 70 μm filter (ThermoFisher Scientific) with excess 1X DPBS (Gibco). Filtrate was subsequently passed through a 40 μm filter (Falcon) with excess 1X DPBS (Gibco). Filtered samples were spun down at 400xg for 5 minutes, resuspended in 1X DPBS, and the equivalent of one quadricep-ECM was plated/well on a 96-well round bottom (Sarstedt).

#### Surface marker staining

Samples were first stained for viability with ZombieNIR Fixable Viability Stain (BioLegend) for 30 minutes on ice in the dark. Samples were washed twice with Fluorescence-Activated Cell Sorting (FACS) buffer, 1X DPBS (Gibco) containing 1% Bovine Serum Albumin (Sigma) and 1 mM EDTA (Invitrogen). Surface marker antibodies were prepared in FACS buffer containing 1:20 TruStain FcX anti-mouse CD16/CD32 (BioLegend), 1:50 Super Bright Complete Staining Buffer (eBioscience), and 1:50 True-Stain Monocyte Blocker (BioLegend) and samples were stained for 45 minutes on ice in the dark. Viability and surface marker antibodies are listed in SI Appendix, Table S2. Samples were washed twice with FACS buffer and fixed with 100 μL FluoroFix buffer (BioLegend) at room temperature for 15 minutes. Samples were washed twice with FACS buffer, resuspended in 1X DPBS, and stored at 4°C overnight. Samples were washed once with 1X DPBS and run on the Aurora flow cytometer (Cytek Biosciences). Gating scheme located in SI Appendix, Fig. S8 was utilized for gating.

#### Intracellular cytokine staining

Samples were stimulated with 200 μL of 1X Cell Stimulation Cocktail (Plus Protein Transport Inhibitors) (eBioscience) in Iscove’s Modified Dulbecco’s Medium (Gibco) containing 5% Fetal Bovine Serum (Gibco) at 37°C for 4 hours. Samples were washed twice in 1X DPBS. Samples were stained with ZombieNIR Fixable Viability Stain (BioLegend) for 30 minutes on ice in the dark. Samples were washed twice with Fluorescence-Activated Cell Sorting (FACS) buffer, 1X DPBS (Gibco) containing 1% Bovine Serum Albumin (Sigma) and 1 mM EDTA (Invitrogen). Surface marker antibodies were prepared in FACS buffer containing 1:20 TruStain FcX anti-mouse CD16/CD32 (BioLegend), 1:50 Super Bright Complete Staining Buffer (eBioscience), and 1:50 True-Stain Monocyte Blocker (BioLegend) for 45 minutes on ice in the dark. Samples were washed twice with FACS buffer and fixed utilizing Cyto-Fast Fix/Perm Buffer Set (BioLegend) according to manufacturer’s instructions. Briefly, samples were fixed with Cyto-Fast Fix/Perm solution (BioLegend) for 20 minutes at room temperature in the dark. 1X Cyto-Fast Perm Wash solution was prepared in deionized water. Intracellular marker antibodies were prepared in 1X Cyto-Fast Perm Wash solution (BioLegend) and samples were stained for 20 minutes at room temperature in the dark. Viability, surface, and cytokine marker antibodies are listed in SI Appendix, Table S3. Samples were washed twice in 1X Cyto-Fast Perm Wash solution (BioLegend), resuspended in 1X DPBS, and stored overnight at 4°C overnight. Samples were washed once with 1X DPBS and run on the Aurora flow cytometer (Cytek Biosciences). Gating scheme located in SI Appendix, Fig. S9 was utilized for gating.

### Histological staining and imaging

#### Tissue preparation

Tissue was formalin-fixed in 10% neutral buffered formalin (Sigma) for 48 hours. Fixated tissue was dehydrated with ethanol, cleared with xylene, and washed and stored in paraffin overnight at 58-60°C. Tissue was embedded in paraffin blocks and sectioned through the Johns Hopkins Oncology Tissue Services SKCCC core facility.

#### Immunofluorescence staining

7 μm formalin-fixed paraffin embedded (FFPE) sections of quadriceps implanted with ECM were backed for 1 hour at 58-60°C. Sections were deparaffinized and rehydrated through sequential steps with xylene, ethanol, and Type 1 H2O. Antigen retrieval was performed in antigen retrieval buffer (AR6, Akoya Biosciences) in steaming conditions for 15 minutes. Sections were cooled to room temperature for approximately 15 minutes and rinsed in Type 1 H2O. Peroxidases were quenched with 3% H2O2 for 15 minutes. Slides were blocked with blocking buffer (1X DPBS (Gibco) containing 10% BSA (Sigma) and 0.05% tween 20 (Fisher Scientific)) for 30 minutes.

Sections were incubated with 1:1000 Rabbit anti-dystrophin antibody (ab218198) for 30 min. Sections were washed three times with 1X Tris Buffered Saline with Tween 20 (TBST) (Cell Signaling Technology) at 3-minute intervals on a shaker. Sections were incubated with the secondary antibody, Rabbit-on-mouse IgG HPR polymer (RMR622H, Biocare Medical). Sections were washed three times with 1X TBST at 3-minute intervals on a shaker and incubated with 1:150 Opal 650 diluted in 1X Amp Diluent for 10 minutes. Sections were washed three times with 1X TBST at 3-minute intervals, stained with DAPI (Akoya Biosciences) for 5 minutes, and rinsed with Type 1 H2O. DAKO mounting medium (Agilent Technologies) was applied and slides cover slipped (0.13-0.16 thickness, FisherScientific). Cover slipped slides were dried overnight at room temperature in the dark and imaged on a Zeiss Axio Imager A2 and stitched with the ZEN software.

#### H&E and Masson’s Trichrome

7 μm formalin-fixed paraffin embedded (FFPE) sections of swiss-rolled colon tissue were stained for H&E by the Johns Hopkins Oncology Tissue Services SKCCC core facility. 7 μm formalin-fixed paraffin embedded (FFPE) sections of quadriceps implanted with ECM were stained for Masson’s Trichrome by the Johns Hopkins Oncology Tissue Services SKCCC core facility. All stained slides were imaged with a digital slide scanner (Hamamatsu NanoZoomer-XR).

#### Bulk RNA Sequencing

Quadriceps implanted with ECM at day 42-post VML surgery were harvested and RNA extracted as described above (See qRT-PCR: Tissue preparation, RNA extraction). Extracted RNA was submitted to Psomagen for QC, Library Prep, and bulk RNA-sequencing. Briefly, TruSeq stranded mRNA library kit was used and pair-end sequencing was performed using the Illumina NovaSeq X platform, generating approximately 50 million paired-end reads per sample. Data was aligned using STAR 2.7.10a against GENCODE GRCm39 vM27. Differential gene expression analysis was performed with DESeq2 v1.42.0 based on a negative binomial model with shrinkage estimation of the logarithmic-fold change (logFC). Gene set enrichment analysis was performed with fgsea v1.28.0, ranking results by the product of logFC and -log10(padj), using REACTOME and HALLMARK pathways. Adjusted p-values were calculated using Benjamini-Hochberg method of multiple testing correction.

### Statistics

All statistical analyses were performed utilizing Graphpad Prism v10 (excluding bulk RNA sequencing analysis). A p-value of less than 0.05 was considered statistically significant. Unpaired two-tailed t-test, one-way ANOVA or two-way ANOVA with appropriate post tests were applied for statistical comparisons, as indicated in the figure legends.

## Acknowledgments

We thank Integra for providing the MicroMatrix UBM Particulate and funding portion of the study. We would to thank Dr. Cynthia Sears, Xinqun Wu, Shaoguang Wu, and Caroline Wensel for technical advice for project planning relating to generating conventionalized germ-free mice and performing 16S rRNA sequencing with Novogene. We would like to thank Hua Ding at the Johns Hopkins Germ-Free Mouse Core for assistance related to all conventionalized germ-free mouse studies. We thank Christopher Cherry and Mike Patatanian for analyzing the bulk RNA sequencing data. We thank Valentine Okechi, Huimin Xi, and Sijia Guo at Novogene for all technical assistance related to project planning, QC, Library Prep, 16S rRNA sequencing, data analysis, and understanding the analysis performed for the 16S rRNA sequencing data set. We thank Hong Wang and Kang Sa at Psomagen for their efforts related to project planning, QC, Library Prep, and bulk RNA sequencing. We thank the Johns Hopkins Oncology Tissue Services SKCCC core facility for sectioning and staining quadriceps and colons used for this study. We would also like to thank Maria Browne for digital slide scanning of the Masson’s and H&E-stained slides. We thank Nathan Price and the animal technicians of the Translational Gerontology Branch at the National Institute on Aging for providing young mouse stool for the conventionalized germ-free study. This study was funded by the NIH Pioneer Award DP1AR076959 (J.H.E.), Bloomberg∼Kimmel Institute for Cancer Immunotherapy (J.H.E, D.M.P.), Bloomberg Philanthropies and the Johns Hopkins University Department of Medicine Chair Fund (C.L.S.), NSF Graduate Research Fellowship Program (B.Y. DGE2139757 and A.R. DGE1746891), NIH Pathway to Independence Award K99AG081564 (J.C.M.), and the Intramural Research Program at the NIA, National Institutes of Health (N.L.P.).

