## Supplementary Materials for "Antibiotic-induced microbiota depletion impairs the pro-regenerative response to a biological scaffold in mice"

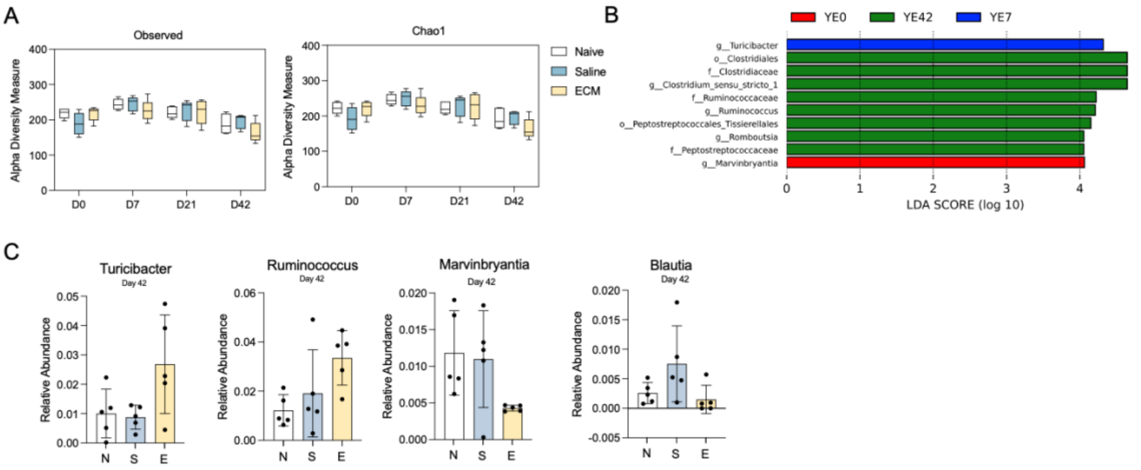


**Fig. S1. Impact of injury and ECM implantation on the diversity and composition of the gut microbiome.** (A) $\alpha$-Diversity of the gut microbiome in naïve, saline, and ECM-treated mice as measured by Observed and Chao1 indices. (B) The linear discriminate effect size (LEFSe) analysis of statistically significant species in ECM implant groups at day 0- (YE0), day 7- (YE7), and day 42-post VML-ECM (YE42). (C) Quantification of the relative abundance of select genus-level bacteria in naïve, saline-treated, and ECM-treated mice over time at day 42-post VML. (A) Ordinary two-way ANOVA with Tukey's multiple comparisons test. (C) Kruskal-Wallis test with Dunn's multiple comparisons test. *P < 0.05, **P < 0.01, ***P < 0.001.

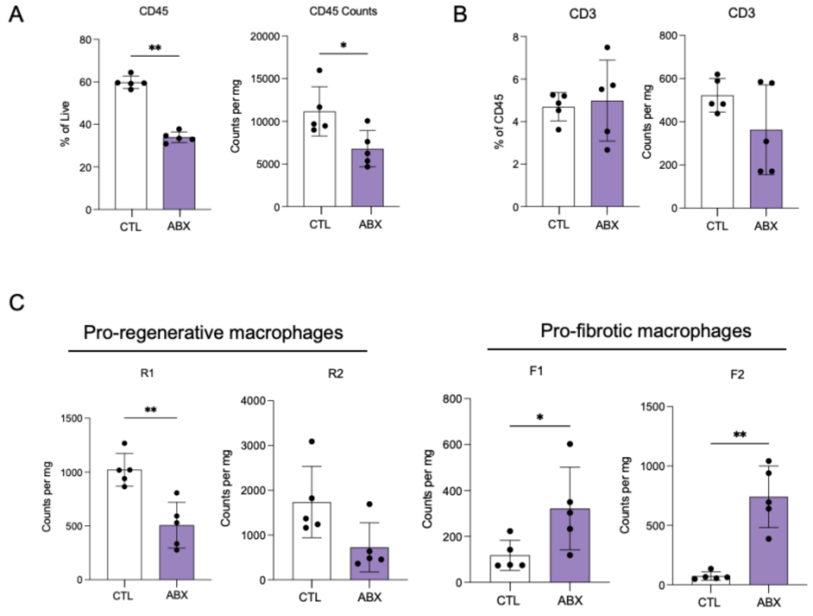


**Figure S2. Impact of antibiotic depletion on immune infiltrate in quadriceps at day 7-post VML-ECM using Pan Immune Flow Cytometry Panel.** (A) Flow cytometric quantification of $CD45^+$ immune cells of ECM-treated muscle at day 7-post VML-ECM. (B) Flow cytometric quantification (frequency and counts per mg of tissue) of $CD3^+$ T cells of ECM-treated muscle at day 7-post VML-ECM (frequency and counts per mg of tissue) (C) Flow cytometric quantification (counts per mg of tissue) of R1, R2, F1, and F2 macrophages of ECM-treated muscle at day 7-post VML-ECM. Bar graphs show mean ± SD. Data are representative of *n* = 5 mice. Mann-Whitney test. *P < 0.05, **P < 0.01, ***P < 0.001.

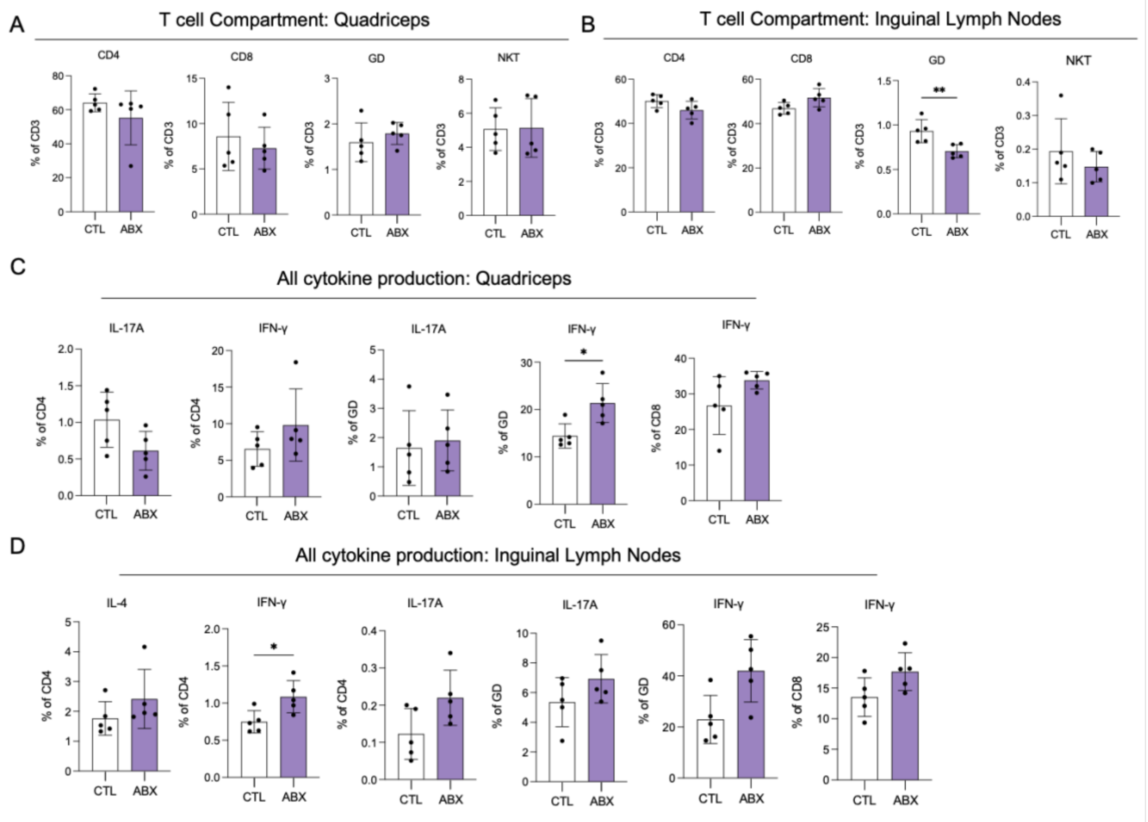


**Figure S3. Impact of antibiotic depletion on all T cell populations and cytokine production in quadriceps and inguinal lymph nodes at day 7-post VML-ECM using the Intracellular Cytokine Flow Cytometry Panel.** (A) Flow cytometric quantification of $CD3^+$ T cell compartment of ECM-treated muscle at day 7-post VML-ECM. (B) Flow cytometric quantification of $CD3^+$ T cell compartment in the inguinal lymph nodes (iLN) of ECM-treated muscle at day 7-post VML-ECM. (C) Flow cytometric quantification of cytokines of ECM-treated muscle at day 7-post VML-ECM. (D) Flow cytometric quantification of cytokines in the inguinal lymph nodes (iLN) of ECM-treated muscle at day 7-post VML-ECM. Bar graphs show mean ± SD. Data are representative of *n* = 5 mice. Mann-Whitney test. *P < 0.05, **P < 0.01, ***P < 0.001.

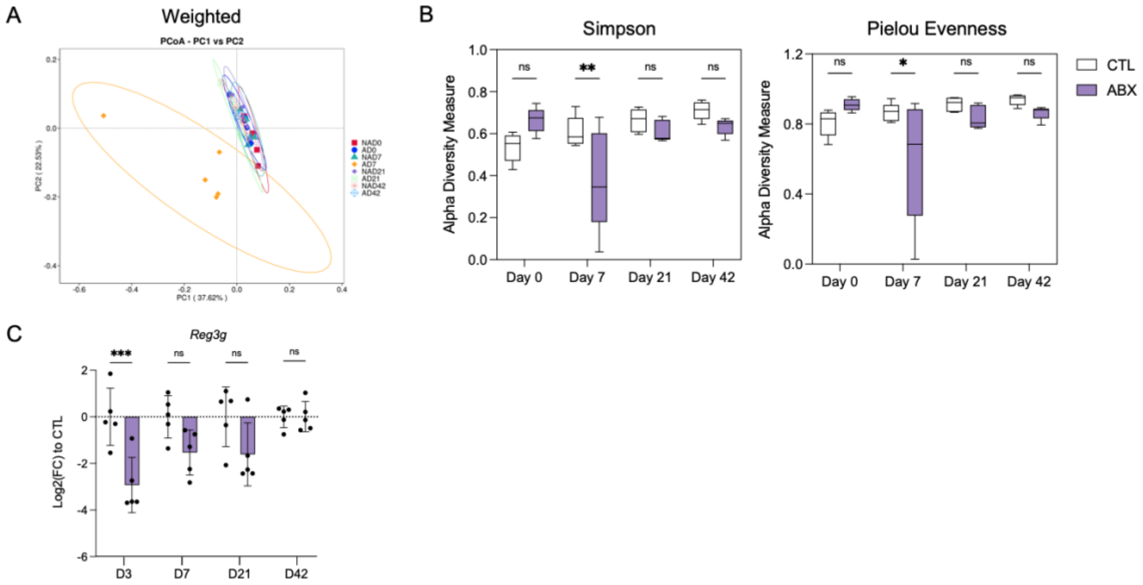


**Figure S4. Assessment of the gut microenvironment in control and antibiotic treated groups.** (A) PCoA based on weighted UniFrac distances. (B) α-Diversity of the gut microbiome in control and antibiotic-treated mice as measured by Simpson's and Simpson's indices and Pielou's Evenness index. (C) Quantification of *Reg3g* gene expression using qRT-PCR in control and antibiotics-treated colon tissue over time.

A

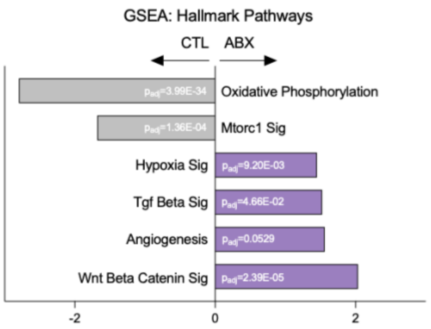


**Figure S5. Impact of antibiotic depletion on transcriptional activity of the quadricep at day 42-post VML-ECM.** (A) Quantification of selected Hallmark pathways from gene set enrichment analysis (GSEA) with normalized enrichment scores (NES).

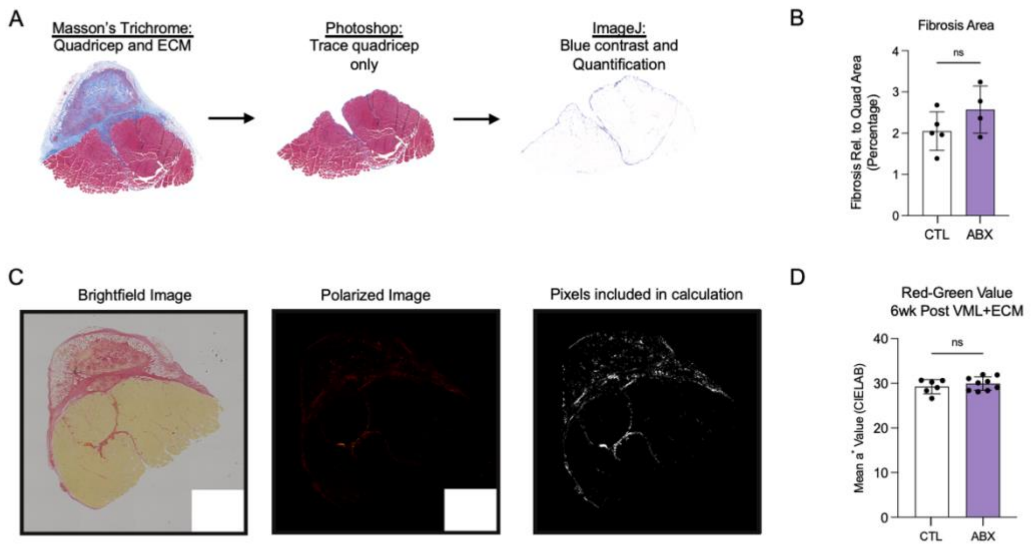


**Figure S6. Evaluation of fibrosis at day 42-post VML-ECM in antibiotics treated mice.** (A) Masson's Trichrome analysis method. (B) Quantification of fibrosis relative to the quadricep area (percentage). (C) Picrosirius Red analysis method. (D) Quantification of mean red-green value on CIELAB axis. Mann-Whitney test. ns=not significant; *P < 0.05, **P < 0.01, ***P < 0.001.

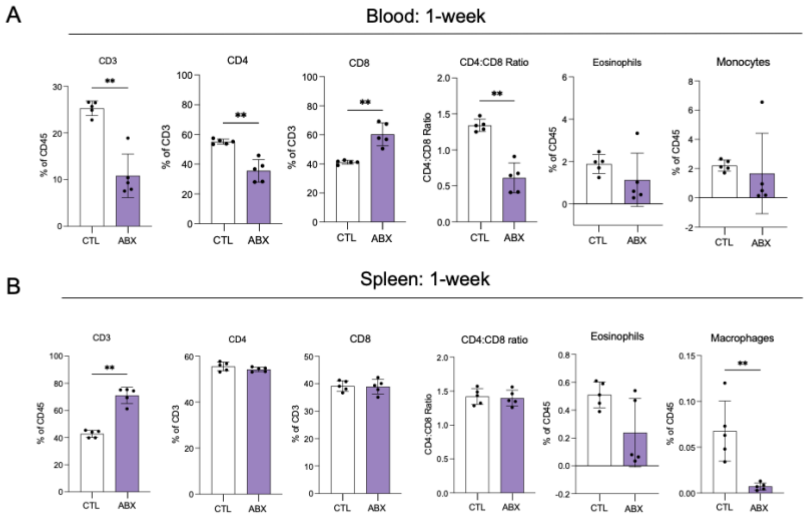


**Figure S7. Impact of antibiotic depletion on immune cell composition in blood and spleen at day 7-post VML-ECM.** (A) Flow cytometry quantification (frequencies) of the indicated immune cell population in the blood of ECM-treated mice at day 7-post VML-ECM. (B) Flow cytometry quantification (frequencies) of the indicated immune cell population in the spleen of ECM-treated mice at day 7-post VML-ECM. Bar graphs show mean ± SD. Data are representative of $n$ = 5 mice. Mann-Whitney test. *P < 0.05, **P < 0.01, ***P < 0.001.

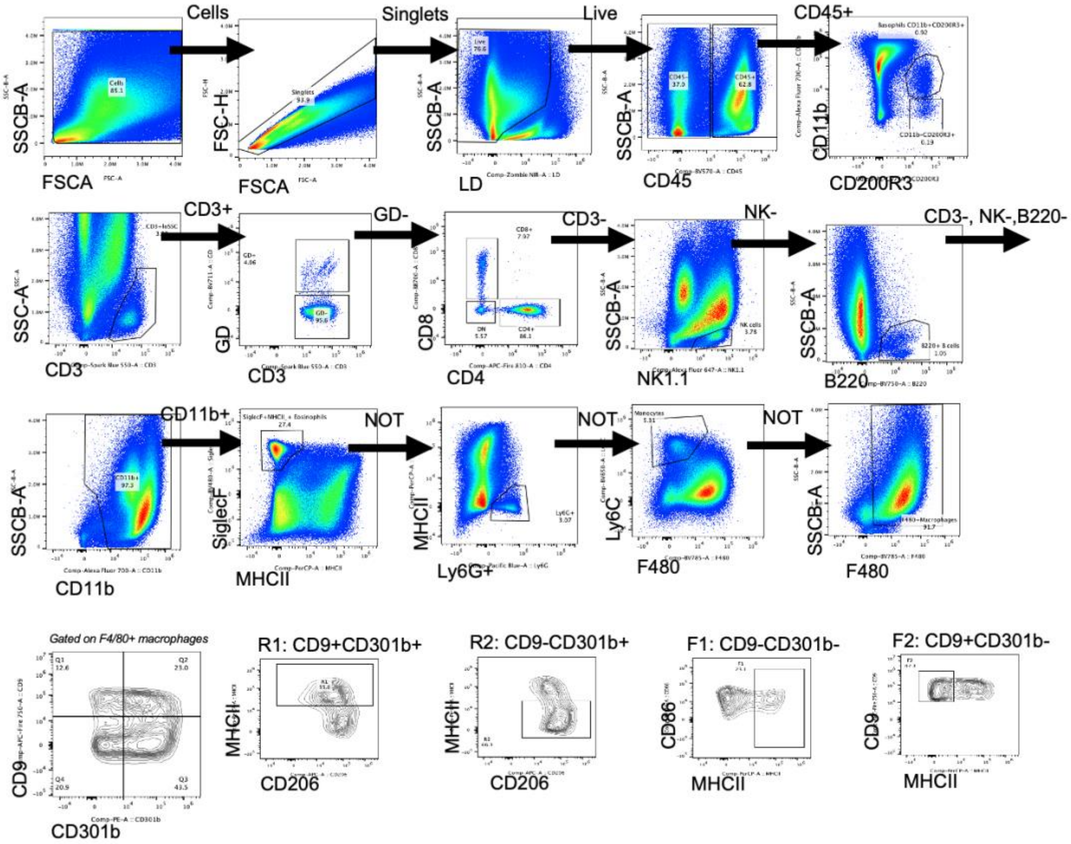


**Figure S8. Gating scheme for Pan Immune flow cytometry panel utilized for VML+ECM quadricep tissue.**

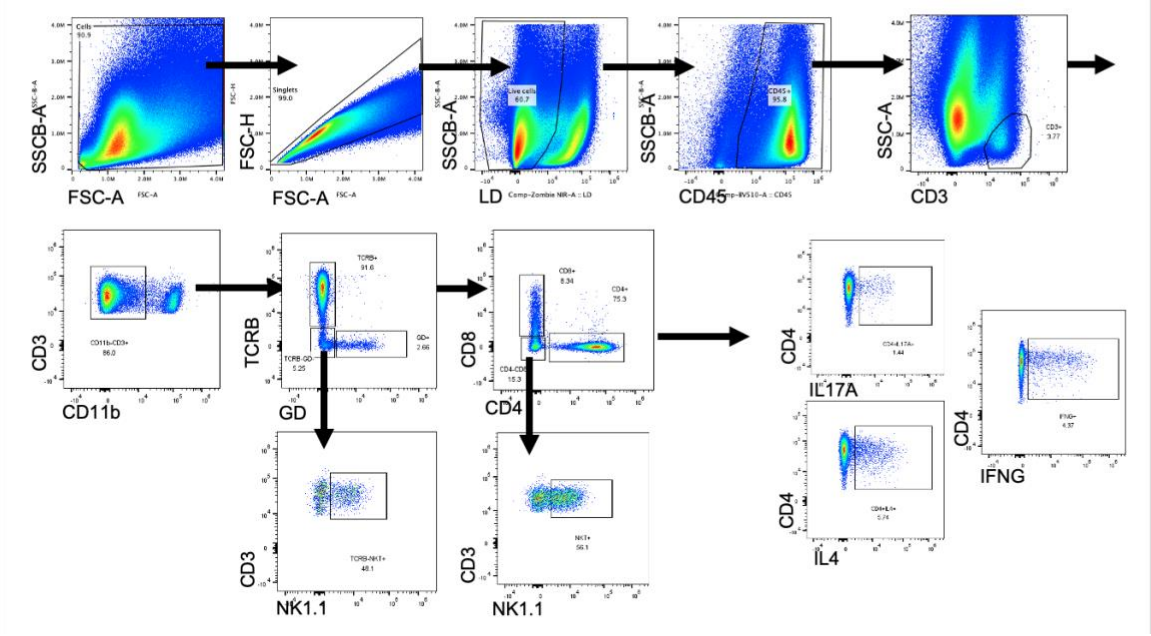


**Figure S9. Gating scheme for Intracellular Cytokine Staining flow cytometry utilized for VML+ECM quadricep tissue.**

**Table S1.** Murine TaqMan gene expression probes.

| Probe: | Assay ID: |
| --- | --- |
| *Gapdh* | Mm99999915_g1 |
| *Il4* | Mm00445259_m1 |
| *Il17f* | Mm00521423_m1 |
| *Ifng* | Mm01168134_m1 |
| *Muc2* | Mm01276676_m1 |
| *Reg3g* | Mm00441127_m1 |
| *Rer1* | Mm00471276_m1 |

**Table S2.** Cytek Aurora Pan Immune Flow Cytometry Panel.

| Fluorophore | Antigen | Clone | Dilution | Manufacturer |
|---|---|---|---|---|
| BV421 | CD86 | GL-1 | 200 | BioLegend |
| SB436 | CD19 | 1D3 | 100 | ThermoFisher |
| Pac Blue | Ly6G | 1A8 | 250 | BioLegend |
| BV480 | SiglecF | E50-2440 | 100 | BD Biosciences |
| BV605 | CD45 | 30-F11 | 300 | BioLegend |
| BV650 | Ly6C | HK1.4 | 1200 | BioLegend |
| BV711 | γδ TCR | GL3 | 200 | BD Biosciences |
| BV785 | F4/80 | BM8 | 300 | BioLegend |
| SparkBlue550 | CD3 | 17A2 | 100 | BioLegend |
| PerCP | MHCII | M5/114.15.2 | 200 | BioLegend |
| BB700 | CD8a | 53-6.7 | 100 | BD Biosciences |
| PE | CD301b | URA-1 | 500 | BioLegend |
| PE-Dazzle594 | CD11c | N418 | 500 | BioLegend |
| APC | CD206 | C068C2 | 200 | BioLegend |
| AF647 | NK1.1 | PK136 | 200 | BioLegend |
| AF700 | CD11b | M1/70 | 400 | BioLegend |
| Zombie NIR | Viability | - | 5000 | BioLegend |
| APC-Fire750 | CD9 | MZ3 | 500 | BioLegend |
| APC-Fire810 | CD4 | GK1.5 | 100 | BioLegend |

**Table S3.** Cytek Aurora Intracellular Cytokine Flow Cytometry Panel.

| Fluorophore | Antigen | Clone | Dilution | Manufacturer |
|---|---|---|---|---|
| BV421 | TCRβ | H57-597 | 100 | BioLegend |
| BV510 | CD45 | 30-F11 | 200 | BioLegend |
| BV605 | NK1.1 | PK136 | 200 | BioLegend |
| BV650 | CD11b | M1/70 | 400 | BioLegend |
| BV711 | γδ TCR | GL3 | 200 | BD Biosciences |
| BV785 | CD4 | GK1.5 | 200 | BioLegend |
| SparkBlue550 | CD3 | 17A2 | 100 | BioLegend |
| BB700 | CD8a | 53-6.7 | 100 | BD Biosciences |
| PE | IL-4 | 11B11 | 100 | BioLegend |
| APC | IFNγ | XMG1.2 | 150 | BioLegend |
| AF700 | IL-17A | TC11-18H10.1 | 200 | BioLegend |
| ZombieNIR | Viability | - | 5000 | BioLegend |
